# Enhanced habit formation in Tourette syndrome: dopamine release and striatal disinhibition modulate shortcut connections in a hierarchical model of cortico-basal ganglia loops

**DOI:** 10.1101/2021.02.08.430235

**Authors:** Carolin Scholl, Javier Baladron, Julien Vitay, Fred H. Hamker

**Affiliations:** Chemnitz University of Technology, Straße der Nationen 62, 09107 Chemnitz, Germany; Max Planck School of Cognition, 04103 Leipzig, Germany

**Keywords:** Tourette syndrome, goal-directed behavior, habitual behavior, basal ganglia, dopamine

## Abstract

In the Gilles de la Tourette syndrome, tics are often considered as habitual responses towards unwanted premonitory urges. Support for the relationship between tics and habits comes from devaluation protocols, which reveal that unmedicated Tourette patients show an increased tendency towards responses to devalued outcomes. We use a neuro-computational model of hierarchically organized cortico-basal ganglia-thalamo-cortical loops to shed more light on enhanced habit formation of Tourette patients. In our model, habitual behavior emerges from cortico-thalamic shortcut connections, where enhanced habit formation can be linked to faster plasticity in the shortcut or to a stronger feedback from the shortcut to the basal ganglia. Irregular activity in such shortcut connections may have different pathophysiological origins. Based on our model, we explore decreased local striatal inhibition, which may correspond to a loss of inhibitory interneurons, and increased dopaminergic modulation of striatal medium spiny neurons as causes for irregular shortcut plasticity or activation. Both lead to higher rates of response towards devalued outcomes in our model, similar to what is observed in Tourette patients. Our results support the view of tics in Tourette syndrome as maladaptive habits. We suggest to reveal more shortcuts between cortico-basal ganglia-thalamo-cortical loops in the human brain and study their potential role in the development of the Tourette syndrome.

## Introduction

Tourette patients repeatedly make movements and sounds that are not entirely voluntary. Such tics are sometimes described as responses towards involuntary premonitory sensations or urges that stop upon tic execution (Brandt et al., 2016; Kwak et al., 2003; Leckman et al., 1993). This view inspired the comparison of tics and habits, which are automatic and fast, yet inflexible responses towards stimuli. In an outcome devaluation paradigm, unmedicated adult Tourette patients with tics indeed relied on habitual rather than goal-directed behavior, more so than healthy control subjects (Delorme et al., 2016), but refer to de Wit et al. (2018) regarding a debate about outcome-devaluation and habits in humans.

Interestingly, habit reversal training is a commonly used cognitive-behavioral therapy method which aims to replace tics by alternative responses (Dutta and Cavanna, 2013). It relies on the idea that tics share key features with habits and thus, may also have a common neural underpinning (Leckman and Riddle, 2000).

The dichotomy between goal-directed and habitual behavior has often been associated with separate cortico-basal ganglia loops through the dorsomedial and dorsolateral striatum (Yin and Knowlton, 2006; Redgrave et al., 2010). However, the directed transition of goal-directed behavior into habits suggests a more integrated and hierarchical organization of these circuits (Balleine et al., 2015; Yin, 2017; Rusu and Pennartz, 2020). Specifically, the habitual system may represent a lower level of the hierarchy than the goal-directed one.

Baladron and Hamker (2020) proposed a new account to habit formation: A model composed of multiple, hierarchically organized cortico-basal ganglia loops suggests that habitual responses emerge from cortico-thalamo-cortical shortcut connections that bypass the longer and slower route through multiple cortico-basal ganglia loops. Habitual learning transfers behavioral control from the cortico-basal ganglia loops to cortico-thalamo-cortical shortcut connections. Shortcut connections may directly connect sensory cortical areas with the thalamus of lower level loops, resulting in a fast excitation of the premotor cortex and the initiation of action. In rats, a site for such shortcuts may be the infralimbic cortex in the medial prefrontal cortex, as lesions of this area prevent them from learning habits (Killcross and Coutureau, 2003); yet when the cortical disruption is applied after the learning of habits, goal-directed behavior reoccurs (Coutureau and Killcross, 2003; Smith et al., 2012). The model of Baladron and Hamker (2020) provides a framework to understand the ineffectiveness of outcome devaluation after overtraining in rodents (Smith and Graybiel, 2013; Adams, 1982): animals keep responding towards devalued outcomes because habitual actions are triggered as direct responses to stimuli via shortcuts, circumventing a careful evaluation of goals.

Here, we aim to investigate whether theoretically grounded changes that simulate the suspected pathophysiology of Tourette syndrome, as described in the next paragraphs, may modulate the effect of shortcut connections and in turn produce the enhanced habit formation observed in the outcome devaluation experiment by Delorme et al. (2016). Specifically, we propose that aberrant activation of cortico-thalamo-cortical shortcut connections may increase the rate of response towards stimuli associated with devalued outcomes, resembling the behavior of Tourette patients in the study.

A first hypothesis of Tourette pathophysiology focuses on disturbed dopaminergic signaling within the basal ganglia (Singer et al., 1982). It has long been assumed that tonic dopamine levels may be reduced in Tourette patients, while in turn phasic dopamine bursts would be increased (Singer et al., 2002; Wong et al., 2008). Yet, because dopamine reuptake inhibitors are less effective in treating tics as expected, Maia and Conceição (2017, 2018) suggest that both tonic *and* phasic dopamine may be increased in Tourette Syndrome. In this framework, increased phasic dopamine bursts may accelerate tic learning by amplifying long-term potentiation on cortico-striatal projections in the direct pathway and long-term depression on projections in the indirect pathway. Increased tonic levels of dopamine may additionally up- and down-regulate the excitability of D1 and D2 striatal cells, thereby reducing the inhibition through the indirect pathway and promoting tic execution (Maia and Conceição, 2017).

A second prominent hypothesis of Tourette pathophysiology involves the feedforward and feedback inhibitory circuits within the striatum. In post-mortem analyses of brains, the caudate nucleus of Tourette patients was found to have smaller volume compared to healthy control subjects (Peterson et al., 1998, 2003), a difference which has been linked to a loss of inhibitory interneurons (Kalanithi et al., 2005; Kataoka et al., 2010). A reduced number of interneurons may give rise to local disinhibition within the striatum (Assous and Tepper, 2019), which, as a final consequence, would decrease the tonic inhibition of the thalamus and release tics. Animal models of tics seem to support this hypothesis: Tic-like movements after striatal disinhibition have been observed in mice and primates (McCairn et al., 2009; Pogorelov et al., 2015). In rats, experimentally induced acute and chronic striatal disinhibition led to acute and chronic tics, respectively (Bronfeld and Bar-Gad, 2013; Vinner et al., 2017).

Based on these hypotheses, we built a spectrum of neuro-computational models that are used to simulate the task-related behavior of control subjects and Tourette patients reported by Delorme et al. (2016). In the experimental study, unmedicated Tourette patients responded more frequently towards stimuli associated with devalued outcomes, indicative of enhanced habit formation, which may relate to tic formation in Tourette syndrome. We successfully replicated this group difference in our simulations by comparing the task performance among different pathological models and a healthy control model whose parameters have been fit to the control group. In particular, we show that aberrant shortcut activation can make the model rely more on habitual behavior. In line with both hypotheses, such aberrant patterns could be indirectly produced by enhanced dopamine modulation or reduced local striatal inhibition.

## Results

### Task description

We tested our neuro-computational model with the task used by Delorme et al. (2016). In this task, participants had to learn stimulus-response-outcome associations (Figure 1A). On each trial, a closed box labeled with a fruit icon was shown to the subject who was then asked to press the right or left button. A correct response was rewarded with points and an image of an open box that contained a fruit. If the subject pressed the wrong key, no points were awarded and an empty box was shown. Six different stimuli were linked to six different outcomes. After learning, Delorme et al. (2016) conducted a cognitive outcome devaluation and stimulus devaluation tests. In the outcome devaluation test, a few of the possible outcome fruits were crossed out with an X, indicating that they would no longer award points. The participant was instructed to only press a key when a still-valued outcome could be obtained. The stimuli devaluation test followed the same procedure, however, at the beginning of each block, some stimuli, instead of outcomes, were crossed out with an X.

**Figure 1:**
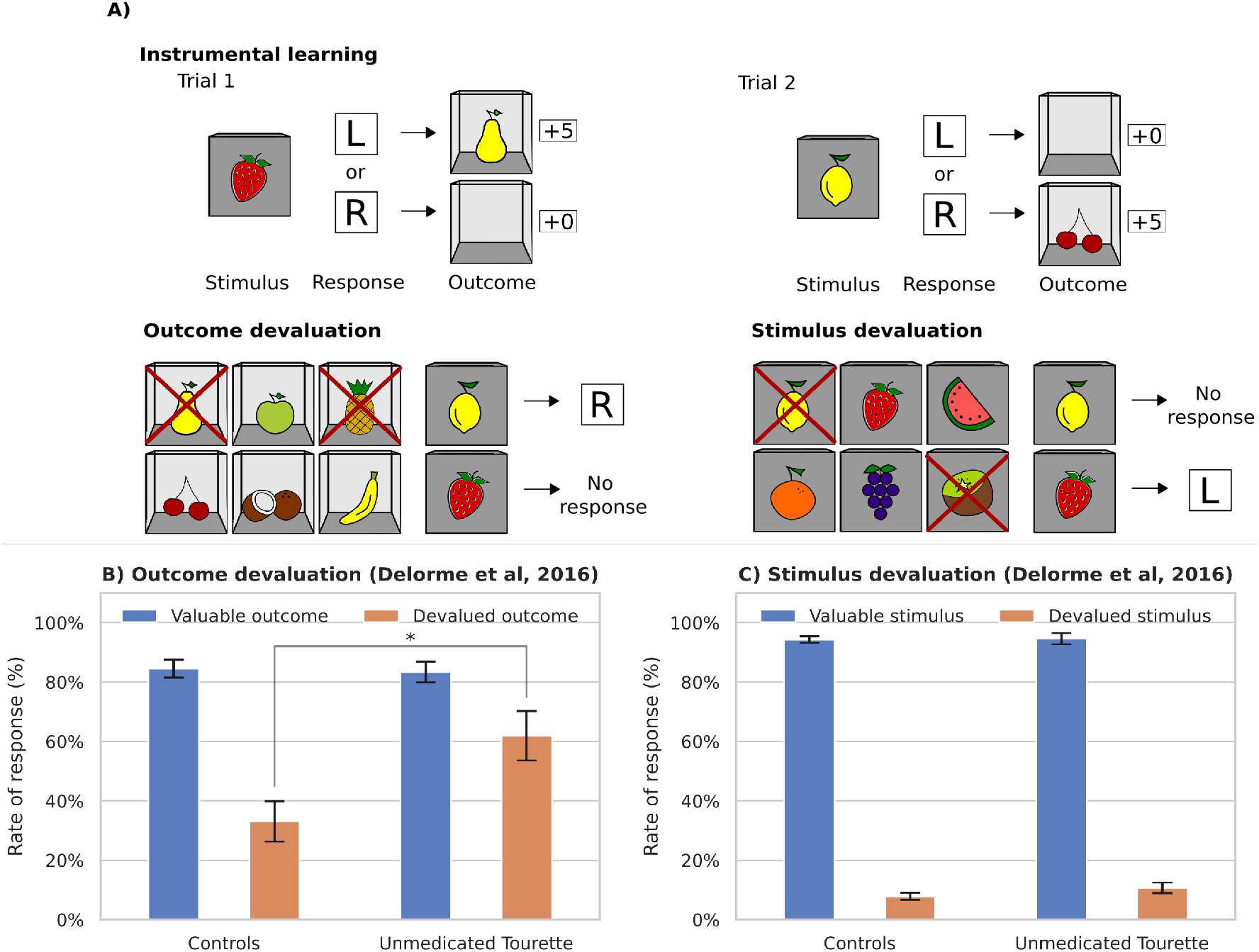
A: Illustration of the task by Delorme et al. (2016). First, participants had to learn associations between 6 stimuli and 6 outcomes by pressing either a left (L) or right (R) button. There was a 100% contingency between stimuli, responses, and outcomes. After successful learning, 2 different outcomes were crossed out per block, marking them as devalued. Participants were instructed to no longer respond to stimuli associated with devalued outcomes. The devaluation of stimuli served as a response inhibition test. Participants were instructed to no longer respond to devalued stimuli. B and C: Experimental results of unmedicated Tourette patients vs. healthy control subjects from the study by Delorme et al. (2016). While there was no difference in the responses towards devalued stimuli between groups (C), unmedicated Tourette patients responded towards stimuli associated with devalued outcomes at a significantly higher rate than healthy controls (B).

Delorme et al. (2016) observed that both, the control group and the Tourette patients, could learn the task. However, during the outcome devaluation test, the patient group presented a higher rate of response towards devalued outcomes (Figure 1B). Further, no significant difference between groups was found in the stimulus devaluation test (Figure 1C). Thus, the difference in the outcome devaluation test could not be attributed to a general deficit in response inhibition of patients.

### Modeling Framework

We adapted the original hierarchical model of multiple cortico-basal ganglia loops of Baladron and Hamker (2020) to simulate the task of Delorme et al. (2016) (Figure 2). The model is composed of a cognitive loop including the dorsomedial striatum and a motor loop including the dorsolateral striatum. Both loops are composed of populations of firing rate units and interact through overlapping cortico-striatal projections (Haber, 2016).

**Figure 2:**
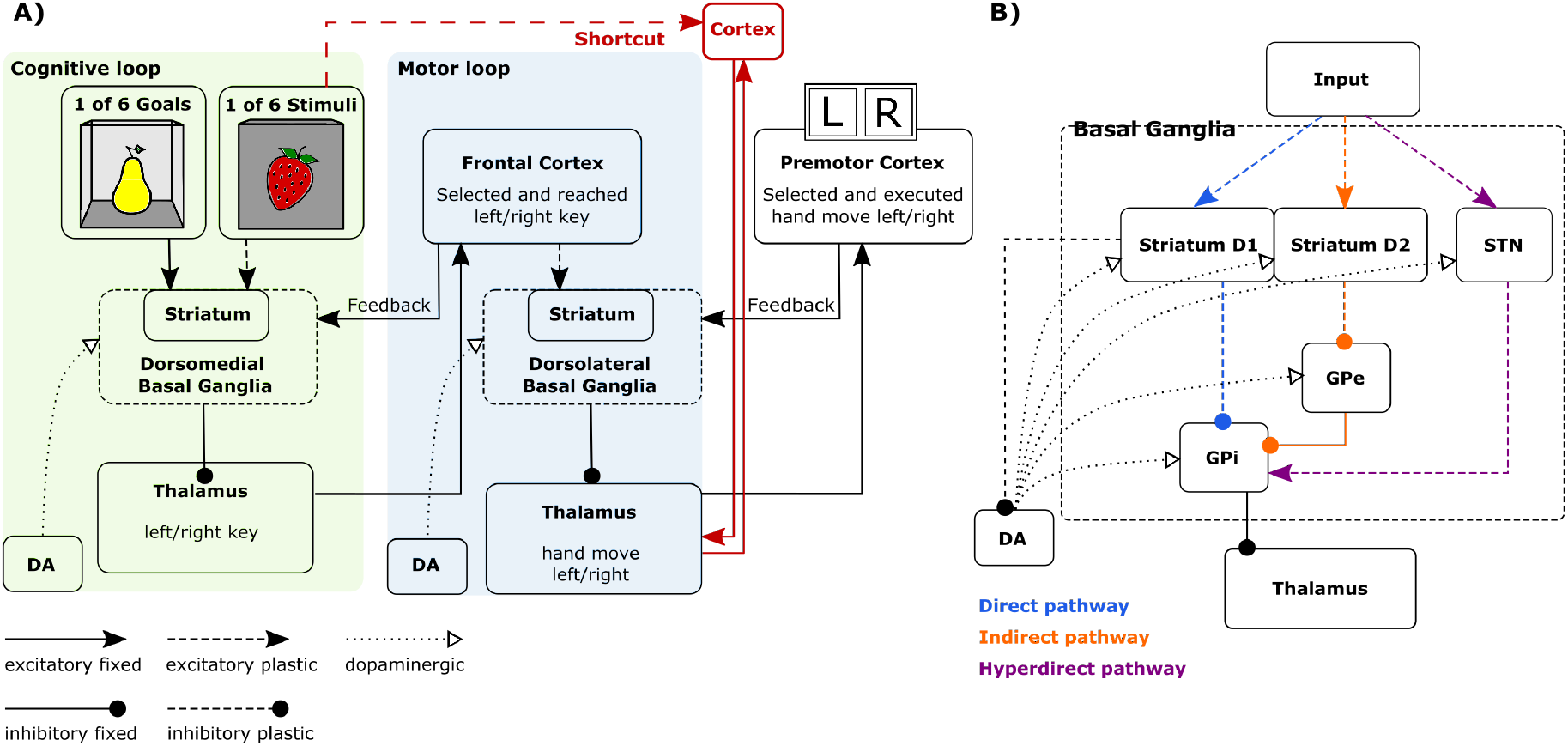
A: Mapping of the task onto the model. In each trial the cognitive loop receives a stimulus signal and an associated goal signal as input, and outputs an objective sent to a premotor loop. A shortcut connects cortical neurons representing the input signal to thalamic cells of the premotor loop. The model response is read from the premotor cortex. B: Each of the two loops includes a direct, indirect, and hyperdirect pathway.

The model of Baladron and Hamker (2020) proposes that an objective, such as obtaining points or food, is divided into a subset of decisions that finally lead to an expected outcome. These decisions are spread between the loops, each of them learning to select an intermediate objective at a different abstraction level, ranging from goals in the ventral striatum to motor commands in the putamen. Each loop provides an objective to the next hierarchical level, which in turn learns to determine the proper decisions to reach it. Further, the model includes cortico-thalamic shortcuts between loops, which are synaptic pathways that can bypass loops. They are trained and monitored by the basal ganglia and they are essential for habitual behaviors.

In the current implementation of the model, the dorsomedial loop receives a desired goal signal (here, a desired outcome fruit) and an associated stimulus signal (observed box with fruit icon) as inputs (Figure 2A). Although the goal selection process is not explicitly modeled, we assume that it involves the limbic network (Groenewegen et al., 1997, 1999; Corbit et al., 2001; Balleine et al., 2003; Gönner et al., 2017), including the ventral striatum (Yael et al., 2019). The dorsomedial loop uses reward signals to learn to select an intended button which is transferred to the dorsolateral loop as a reference signal. The dorsolateral loop then learns to select the appropriate hand movement in the premotor cortex. The model therefore separates the prediction of a state where the desired reward could be obtained from the action required to reach it. Such an organization provides multiple computational benefits such as transferring knowledge between tasks or simplifying the credit assignment problem (Baladron and Hamker, 2020).

Cortico-thalamo-cortical pathways (Sherman and Guillery, 2011), which in our model serve as a shortcut by bypassing the dorsomedial loop, directly link sensory information via cortical representations with hand movements (Figure 2A). This shortcut is monitored and trained in a Hebbian manner by the basal ganglia through its output projections. In our previous work, we have shown how such a pathway can explain the emergence of habitual behavior and the ineffectiveness of outcome devaluation after overtraining (Baladron and Hamker, 2020), as well as the effect of pallidotomy in Parkinsonian patients (Baladron and Hamker, 2015; Schroll et al., 2014).

### Role of cortico-thalamic shortcuts

The shortcut connection links stimuli to the premotor loop (Figure 2A). In order to test the role of the shortcut, we ran experiments with different learning speeds in the shortcut connections, so that the weights changed by a different amount after each co-activation of the presynaptic and postsynaptic cells. One additional set of simulations was performed in which learning in the shortcut was fully disabled. As can be seen in Figure 3A, all models initially select actions randomly, then gradually increase their performance, reaching a value higher than 90% after 8 blocks. This compares well with the results from Delorme et al. (2016), where both patients and controls reached a performance above 90% after the same number of trials.

**Figure 3:**
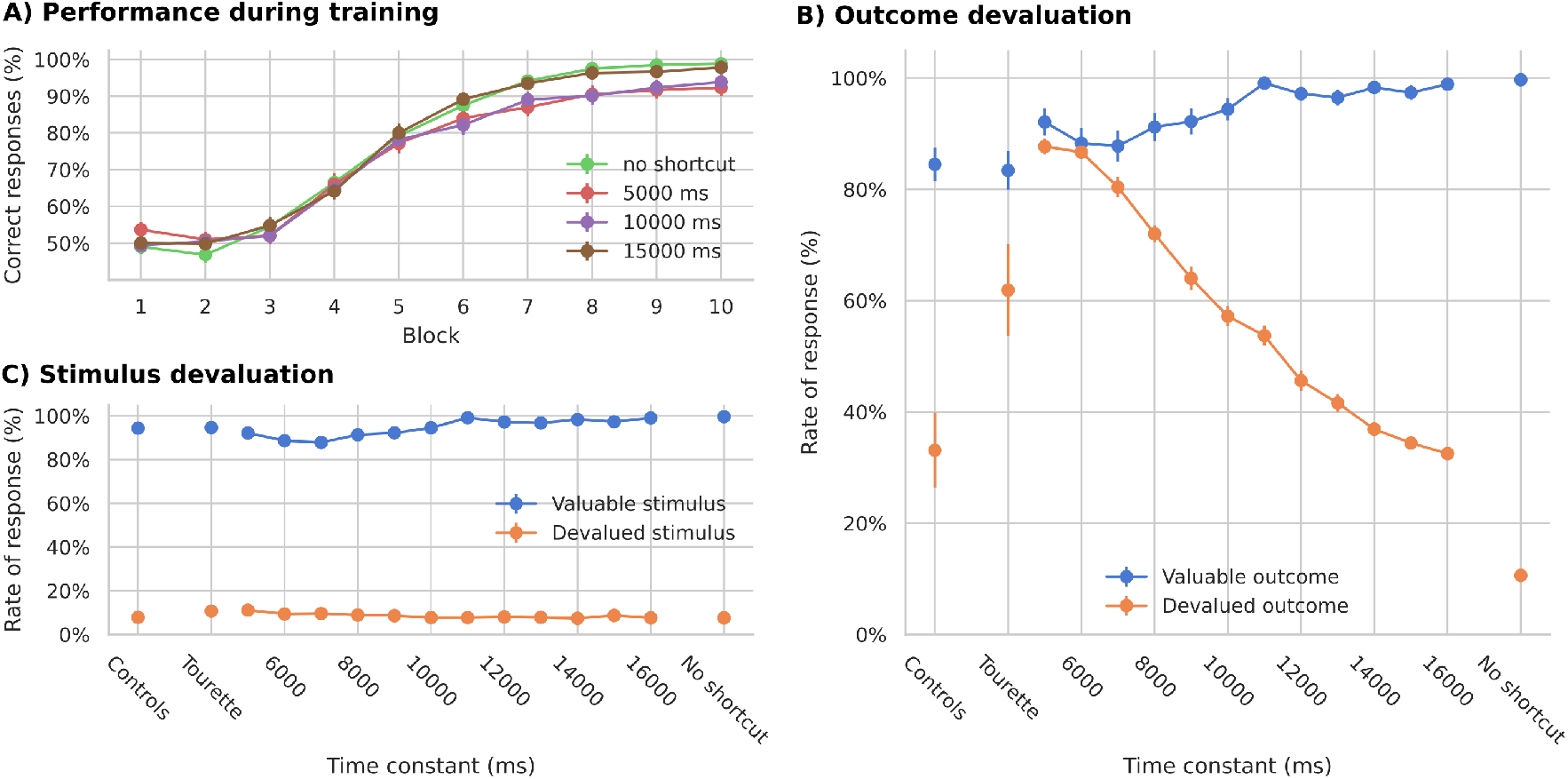
Simulated learning of the action-outcome associations of models with different time constants for learning the shortcut connections. A: Learning performance over 10 blocks of 12 trials each. B: Rate of response towards stimuli associated with devalued outcomes compared to still valuable outcomes. A smaller time constant translates to faster learning. C: Rate of response towards devalued stimuli compared to still valuable stimuli.

Analogous to the experiments from Delorme et al. (2016), we then simulated outcome and stimulus devaluation. Outcome devaluation was simulated by canceling the goal signal reaching the dorsomedial loop on trials associated with a devalued outcome. This represents a lack of interest in the possible reward. Stimulus devaluation was simulated by reducing the input stimulus signal on trials associated with a devalued stimulus. This represents the fact that the stimulus is still observed, but not attended. Two of all possible outcomes or stimuli were devalued in each of the 6 blocks.

Although all versions of the models show similar performance during training (Figure 3A), outcome devaluation has different effects. Models with faster learning in the shortcut (smaller time constant, *τ_w_* in Eq. 13) select devalued outcomes more frequently than models with medium or slow learning speeds (Figure 3B). The results on the stimulus devaluation test however show no significant difference between the models (Figure 3C).

The difference between models with different learning speeds compares well to observations by Delorme et al. (2016) between Tourette patients and controls. On outcome devalued trials, patients had a significantly higher rate of response towards devalued outcomes, suggesting that patients relied more on habitual behavior than control subjects. Further, there was no significant difference between groups in the stimulus devaluation test.

Our results suggest that faster learning in the cortico-thalamic shortcut can explain the difference between controls and patients in the task of Delorme et al. (2016). As in our previous experiments (Baladron and Hamker, 2020), the development of habitual behavior relies on these connections, which bypass the goal analysis done by the dorsomedial loop. To further test this hypothesis, we ran an additional set of simulations in which learning in the shortcut was completely removed. Models without plasticity in the cortico-thalamic shortcut are unable to develop habits and therefore have a much lower response rate to devalued outcomes (Figure 3B). However, their performance on learning the task is similar to that of models with shortcut learning. This confirms the habit learning framework of Baladron and Hamker (2020): habits emerge by learning shortcuts.

In summary, an increase in the shortcut learning speed can drive the model to rely more on habitual behavior, providing a possible explanation to the enhanced habit formation in Tourette patients. Such a change in the model could however emerge from different pathological causes, such as by a direct modification of the shortcut connections or indirectly through abnormal modulation of the components of the learning rule. In the following, we explore whether such an abnormal learning may arise from hypothesized pathophysiologies of Tourette syndrome.

### Enhanced Dopaminergic Modulation

It has been suggested that tics in Tourette syndrome are caused by dopaminergic dysfunction. While the exact anomaly is still debated, theories tend to link higher concentrations of dopamine in the axon terminals to the symptoms (Buse et al., 2013). Following these results, increased habitual responses may be produced due to a strengthened direct pathway and not necessarily through abnormal shortcuts as we propose here. However, in the context of our model, altered dopamine signaling can indirectly affect the shortcut’s behavior by either changing the output of the basal ganglia that trains the shortcut or by modulating the shortcut’s feedback to the striatum, creating a bias towards the action selected by the shortcut.

Dopamine is known to have two different effects on striatal cells (Gerfen and Surmeier, 2011). First, it modulates the activation of cells depending on the dopamine receptor being stimulated (Surmeier et al., 2007). D1 receptor signaling increases the activation while D2 receptor signaling decreases it. Second, dopamine regulates plasticity (Wickens, 2009). An increase in the dopamine level enhances long-term potentiation in cells expressing D1 receptors and long-term depression in cells expressing D2 receptors (Shen et al., 2008; Fisher et al., 2017). We investigate both mechanisms separately in our model to better understand the effect of each on the habitual responses of Tourette patients.

#### Effects of altered response modulation

We first modeled the effect of dopamine on the firing rate of striatal cells. Increased dopamine levels were implemented by introducing a scaling factor from the membrane potential to the firing rate (*S_f_* in Equation 1, Figure 4A). With higher levels of tonic dopamine, the excitability of cells in the direct pathway is increased, while it is decreased in the indirect pathway. The parameters of the plasticity rule were unaffected and the time constant was set to 15,000 ms (Figure 3B).

**Figure 4:**
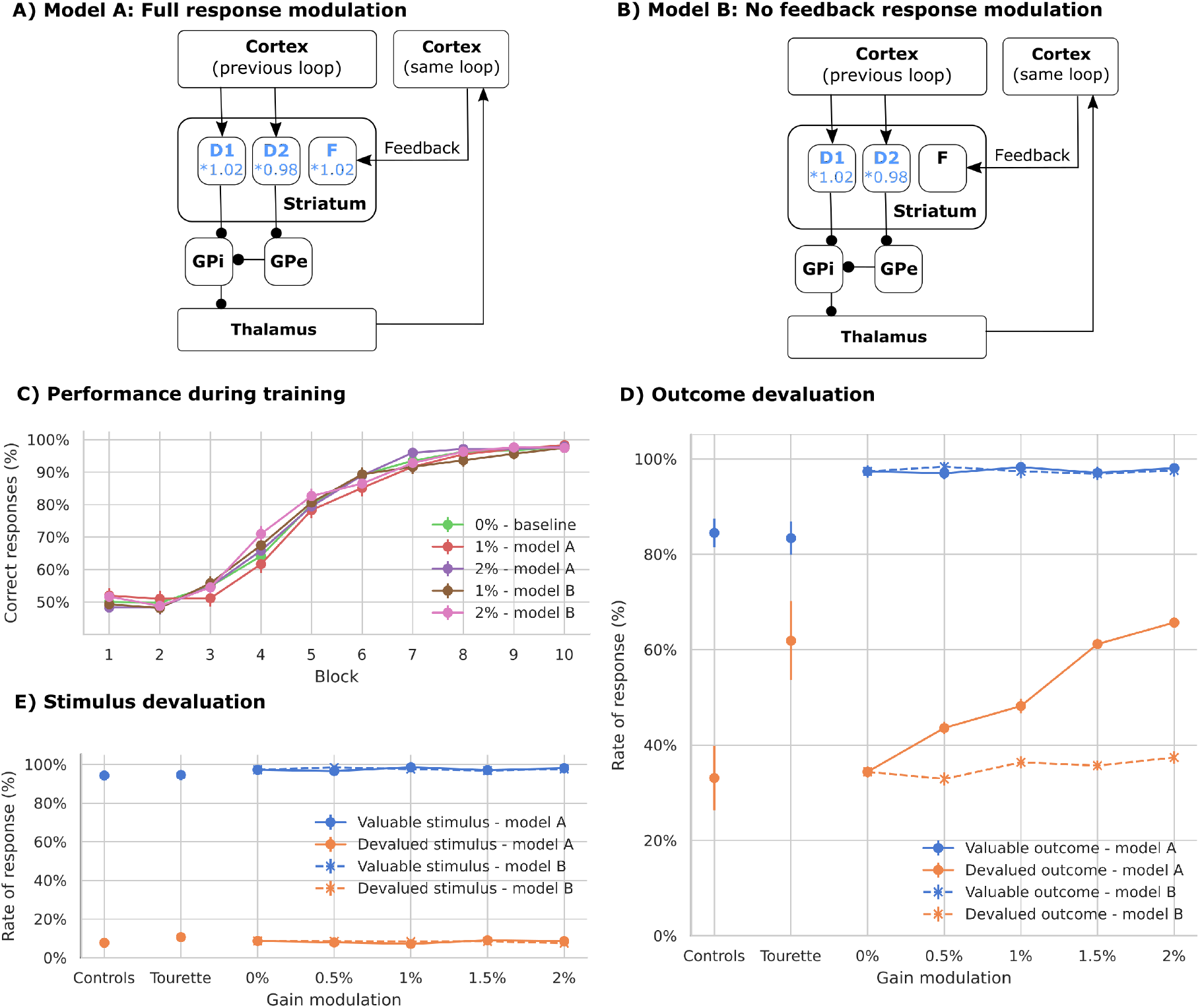
Simulated learning of action-outcome associations of models with dopamine-mediated response modulation of striatal firing rates. All other connections, including the pathway via the STN are modeled as before. A: To test the dopamine-dependent rate modulation hypothesis, the output rate of striatal cells is multiplied by a fixed factor (model A). B: In model B, the response modulation is not applied on the feedback connections. C: Learning performance over 10 blocks of 12 trials each. All models show a similar performance. D: Responses towards stimuli associated with still valuable and devalued outcomes. Only models with full response modulation, including the feedback pathway, show an increased rate of responses. E: Responses towards valuable and devalued stimuli.

There are two possible ways in which dopamine-based response modulation can affect habitual responses in the model. The first option is that the imbalance of the indirect and direct pathways disrupts the outputs of the basal ganglia to the thalamus, which are used to train the shortcut. The second option is that the feedback to the striatum is increased, thereby biasing the response. In order to disentangle the effects, we performed an additional experiment in which we removed the response modulation on those cells receiving cortical feedback (Figure 4B). These feedback projections transmit shortcut activation to the dorsolateral loop and enhance it through the direct pathway. In this condition, response modulation is only present for those cells that receive exclusively inputs from the previous loop through overlapping cortico-striatal projections.

Both options of the model with dopamine-dependent response modulation as well as the control model learn the task and reach similar performance levels (Figure 4C). The models however vary in the rate of responses to devalued outcomes (Figure 4D): A dopamine-dependent modulation of the firing rate leads to more responses to devalued outcomes (significant difference between modulation level 1.02 and control, permutation test p<0.005), similar to those of unmedicated Tourette patients as reported by Delorme et al. (2016). However, this effect depends on the dopamine-dependent modulation of the feedback signal, as models without such a modulation do not increase their responses to devalued outcomes (Figure 4D, no significant difference, p>0.05), while not affecting the learning of the task (Figure 4C).

Shortcut connections can therefore make the model rely more on habitual behavior, not only when the speed of plasticity in shortcuts is increased, but also when the impact of the shortcut on the basal ganglia circuits is increased through a dopamine-dependent modulation of striatal activity.

#### Effects of altered plasticity modulation

Second, we tested whether dopamine modulation of cortico-striatal plasticity could also indirectly affect the shortcut and increase habitual responses. Thus, we increased the impact of dopamine in the learning period after reward delivery, thereby amplifying long-term potentiation in D1 cells and long-term depression in D2 cells (Figure 5A).

**Figure 5:**
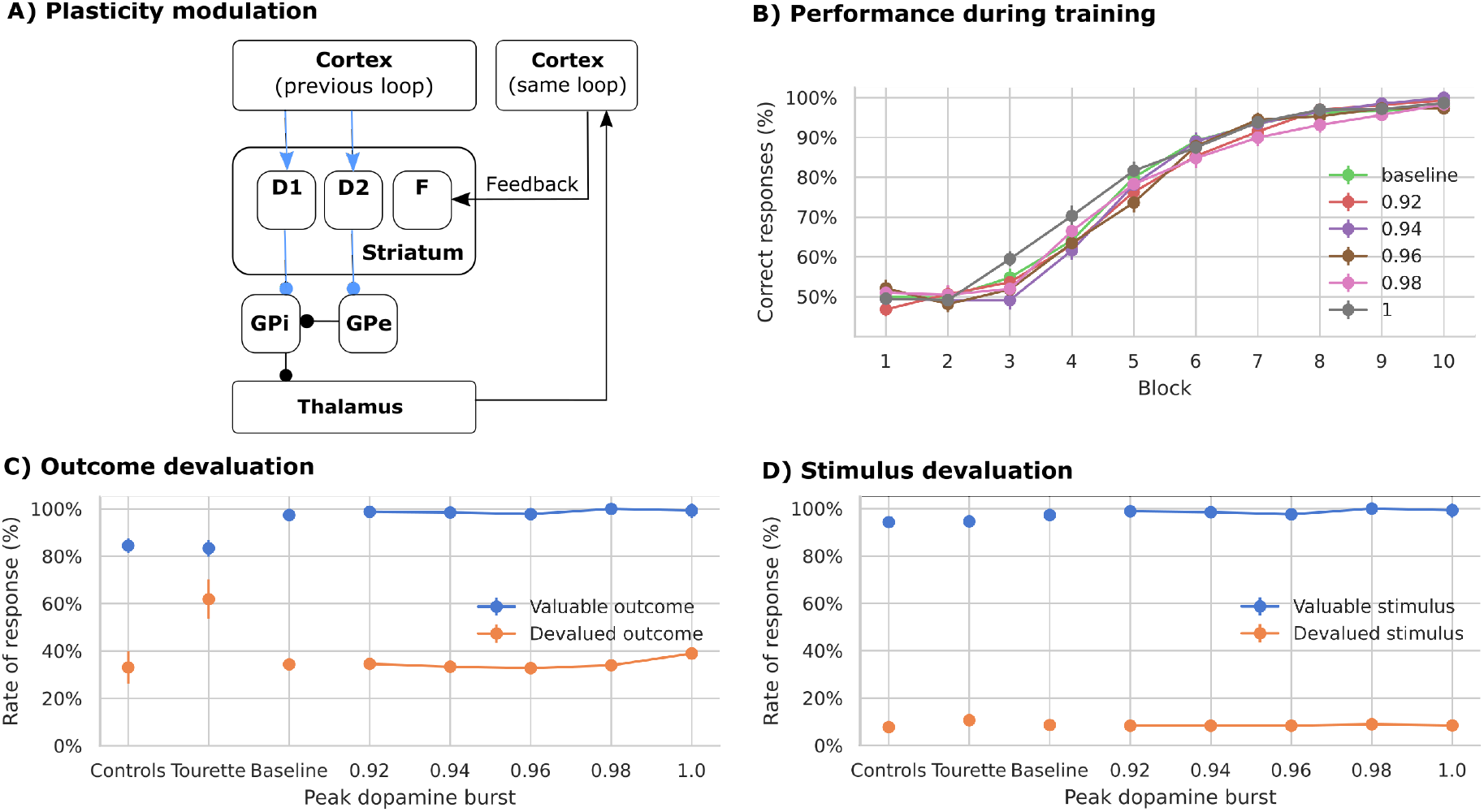
Simulated learning of action-outcome associations of models with dopamine-mediated up- and down-regulation of plasticity of cortico-striatal synapses. A: The effect of the dopamine signal on plasticity is enhanced. In each model, the size of the phasic increase in the dopamine signal after receiving reward is different. The baseline dopamine used to compute the level of a phasic change is varied (see Equation 2). Lowering the baseline increases the maximum amplitude reached by the phasic response. B: Learning performance over 10 blocks of 12 trials each. All models present similar performance. C: Responses towards stimuli associated with still valuable and devalued outcomes. Increasing the amplitude of the dopamine bursts does not significantly change the learning behavior. D: Responses towards valuable and devalued stimuli.

Models with increased plasticity modulation also learn the task (Figure 5B), but their response rate towards devalued outcomes is similar to control models (Figure 5C, no significant difference, p>0.05). Further, all versions show a small and similar amount of responses to devalued stimuli (Figure 5D, no significant difference between the groups).

Thus, our model of increased dopamine-mediated up- and down-regulation of plasticity does not lead to pronounced responses towards devalued outcomes.

### Reduced local inhibition in the striatum

The second major hypothesis regarding Tourette pathophysiology involves reduced striatal inhibition (Bronfeld and Bar-Gad, 2013; Vinner et al., 2017). We therefore tested whether reducing the weights between striatal inhibitory projection neurons would produce any change in the rate of responses to devalued outcomes and could account for the observations of Delorme et al. (2016).

#### Effects of reduced inhibition in the dorsomedial striatum

The performance of the model with lowered inhibitory connections in the dorsomedial striatum is robust to a decrease in the weights down to 40% of the original level (Figure 6A). Models with no local inhibition become unstable at block 5 during learning and do not learn the task well. Models with reduced weights show a trend towards an increased rate of responses towards devalued outcomes (Figure 6B, significant difference, p<0.005). However, the response rate is lower than for patients tested by Delorme et al. (2016). The rate of response to devalued stimuli is similar in all cases (Figure 6C, no significant difference between the groups, p>0.05).

**Figure 6:**
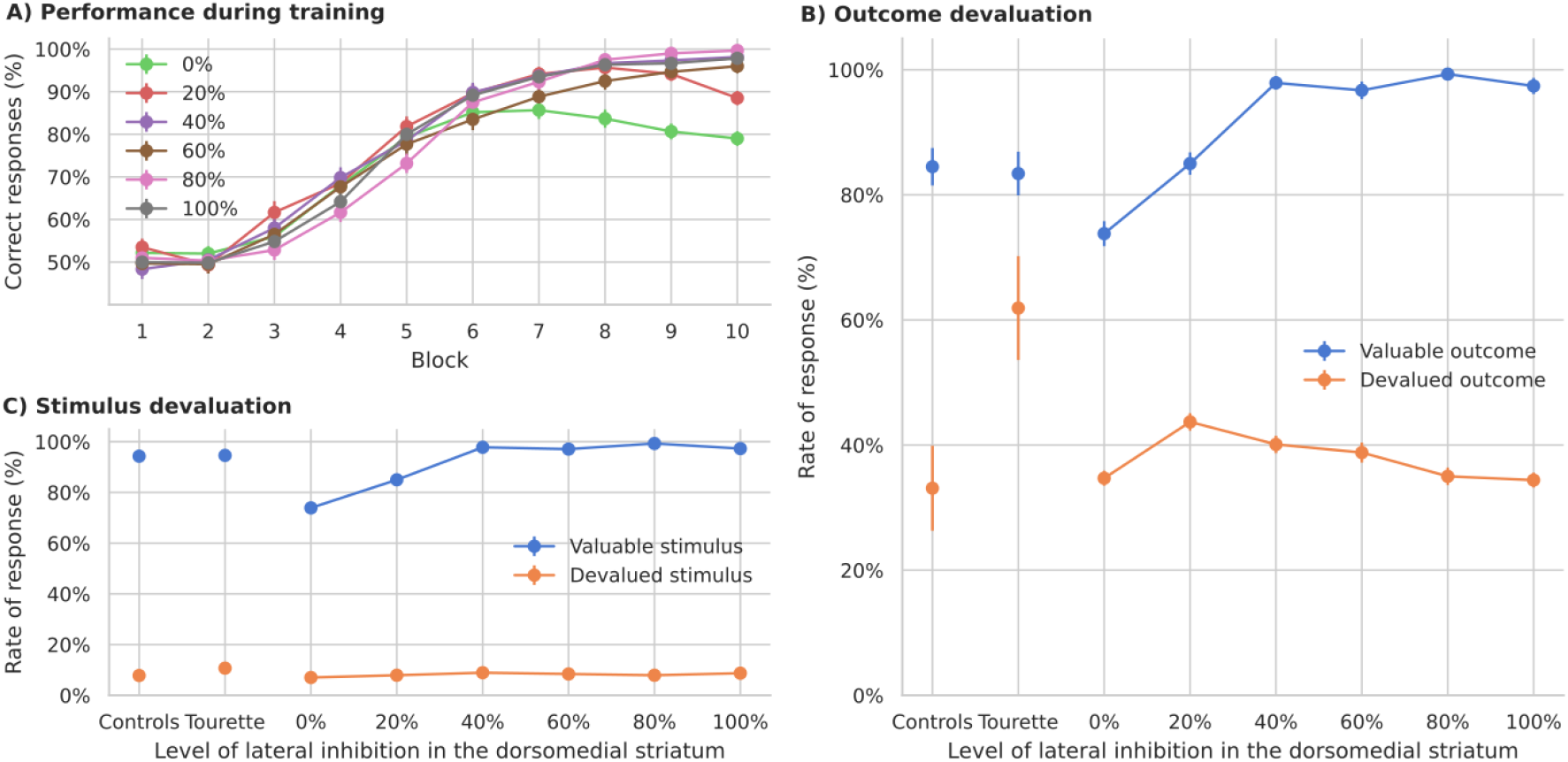
Simulated learning of action-outcome associations of models with reduced local striatal inhibition in the dorsomedial loop. A: Learning performance over 10 blocks of 12 trials each. In each model the weight is reduced by a fraction of their original value (from no inhibition, 0%, to control, 100%). When the inhibition is completely removed, the model becomes unstable. B: Responses towards stimuli associated with valuable and devalued outcomes. The rate of response to devalued outcomes shows a small increase when inhibition is strongly reduced. C: Responses towards valuable and devalued stimuli. Models with a strong reduction show less frequent responses to valuable stimuli.

Models with a strong reduction of dorsomedial striatal inhibition show a higher variability in the weight matrix learned by the shortcut during the task. Although the mean weight value in both the control models (100% inhibition) and those with weights decreased to 40% is the same (0.65), a significant difference occurs in their standard deviation (difference of 0.001, p<0.005). This indicates that models with reduced inhibition produce a variability in shortcut strength that can make them more dependent on habitual behavior. Increased shortcut variability changes the balance in the baseline of the thalamus, allowing the basal ganglia to take over the control through its inhibitory projections. Therefore, unlike the effect of firing rate modulation, reduced inhibition can affect habitual responding by modulating shortcut plasticity directly and not via the feedback connection. Its overall impact however is much smaller.

#### Effects of reduced inhibition in the dorsolateral striatum

We also reduced the inhibition in the striatum of the dorsolateral loop following the same procedure as for the previous loop. This reduction has a stronger effect on the learning performance of the model (Figure 7A) and only those models with a slight reduction of inhibition reach a performance similar to control models and human subjects. Reduced inhibition in the dorsolateral striatum, however, does not lead to an increase in response rate to devalued outcomes comparable to Tourette patients in Delorme et al. (2016) (Figure 7B).

**Figure 7:**
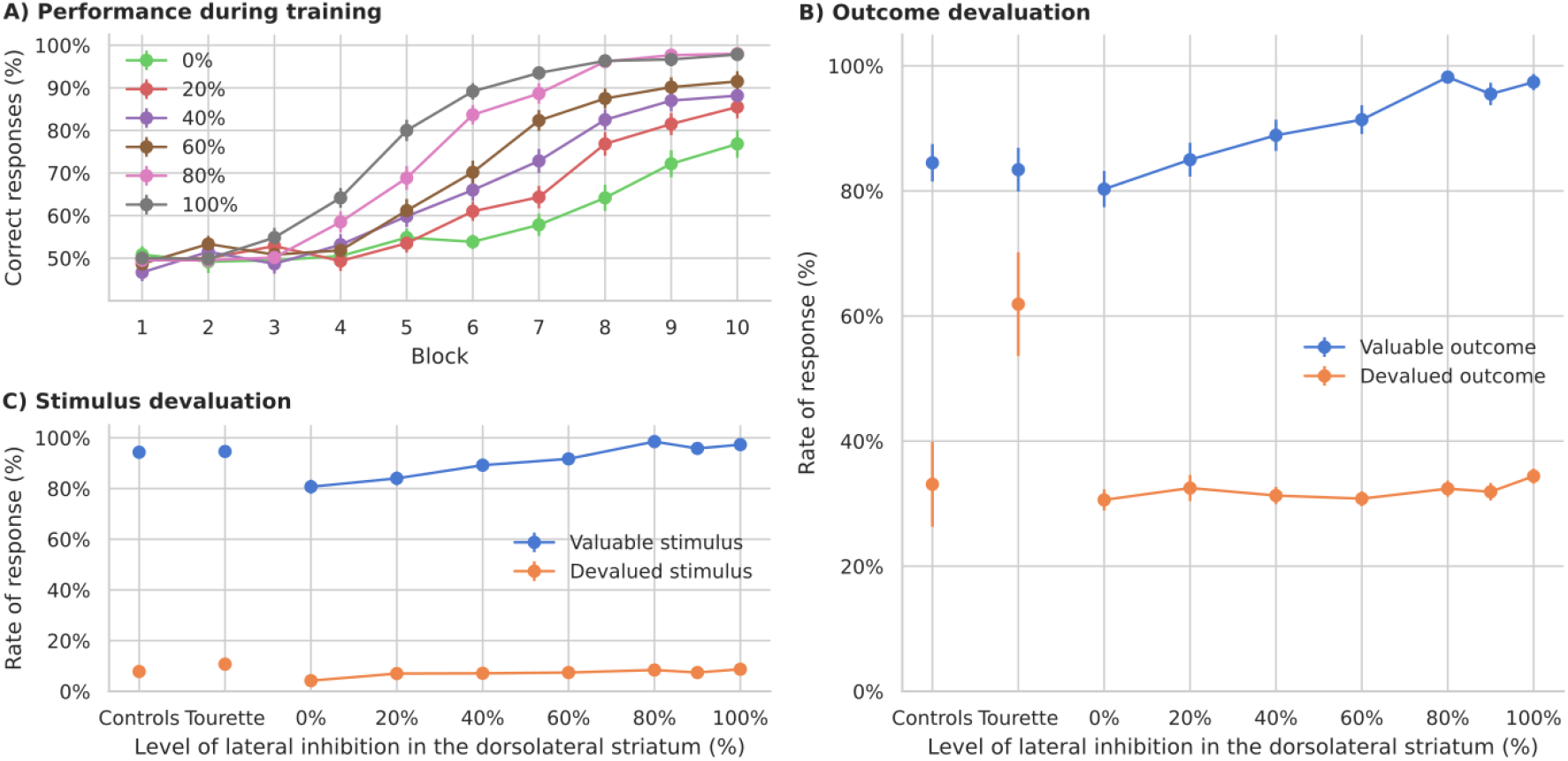
Simulated learning of action-outcome associations of models with reduced local striatal inhibition in the dorsolateral loop. A: Learning performance over 10 blocks of 12 trials each. Only models with a small reduction in inhibition show a performance similar to controls. B: Responses towards stimuli associated with valuable and devalued outcomes. Different levels of local striatal inhibition in the dorsolateral loop do not affect the rate of response to devalued outcomes. C: Responses towards valuable and devalued stimuli. Models with a strong reduction show less frequent responses to both, valuable and devalued stimuli.

Consequently, our model predicts only a modest increase in habitual behavior with reductions of lateral inhibition in the dorsomedial striatum, but no effect to changing inhibition levels in the dorsolateral striatum.

## Discussion

The neurobiological underpinnings of Tourette syndrome are still not completely clear. Two main lines of research focus on explaining tic generation through either imbalanced inhibition (Kalanithi et al., 2005; Kataoka et al., 2010; Vinner et al., 2017) or on anomalies of dopamine transmission (Maia and Conceição, 2018; Maia and Conceição, 2017). Based on the suggested commonalities of tics and habits, we propose a neurocognitive model of enhanced habit formation in Tourette syndrome. The model’s difference in behavior under pathological and default parameter configurations offers new avenues to understand Tourette pathophysiology and complements traditional views.

In our model, two hierarchically organized cortico-basal-ganglia-thalamo cortical loops simulate the increased engagement of habitual behavior by Tourette patients. Sensory inputs can either drive neurons in the dorsomedial striatum directly or, via a cortico-thalamic shortcut, reach the dorsolateral striatum. We here propose that enhanced habit formation in Tourette patients, as observed by Delorme et al. (2016), may be grounded in aberrant activation within cortico-thalamic shortcut connections. Models with faster learning in the shortcut produce similar data as Tourette patients. To better link aberrant learning with the potential pathophysiology of Tourette, we investigated two hypotheses – enhanced dopamine signaling and striatal disinhibition. Enhanced dopamine signaling modulates the activation of the shortcut through the closed loop formed by the direct pathway and feedback cortical connections. Reduced striatal inhibition introduces a high variability in the shortcut. Both changes increase the amount of habitual responses mimicking the behavior of Tourette patients reported by Delorme et al. (2016).

### Relation between Tic Formation and Habit Formation

Cognitive symptoms other than tic generation have been reported in Tourette syndrome (Brand et al., 2002; Eddy and Cavanna, 2013; Puts et al., 2015), but barely discussed. In this work, we primarily address enhanced habit formation of Tourette patients in a cognitive task, and not necessarily tic generation. Yet as our model simulations have shown, altered shortcut behavior can be indirectly produced through the two anomalies that have been discussed in the context of tic formation as well.

Tics have been compared to habits from both a cognitive-behavioral and neuroscientific perspective (Leckman and Riddle, 2000; Maia and Conceição, 2017; Delorme et al., 2016; Shephard et al., 2019; Beste and Münchau, 2018). Our model appears consistent with the cognitive framework of altered perception-action binding in Tourette patients (Beste and Münchau, 2018), as the shortcut binds perceptual states to motor actions via the basal ganglia. Kleimaker et al. (2020) demonstrated that Tourette patients show an increased perception-action binding. This may be regarded as a surplus of actions (Beste and Münchau, 2018), some of which become habits due to an increased propensity towards habit learning (Delorme et al., 2016) and reward learning (Palminteri et al., 2009, 2011). Specifically, tics may be habitual responses towards unpleasant somatosensory internal stimuli, the so-called premonitory urges. The termination of the urge through tic execution can be viewed as avoiding punishment, making future tic execution more likely (Brandt et al., 2016; Capriotti et al., 2014; Kwak et al., 2003), and after repeated execution automatic and habitual. Yet it should be noted that the typical sequence of symptom onset is conflicting with this view. Children typically first report about urges around three years after tic onset (Openneer et al., 2019). This could however also be attributed to missing awareness for urges and unreliable reporting. Finally, habit reversal training is a promising therapy option that views tics as maladaptive habits (Dutta and Cavanna, 2013): it tries to identify the preceding urge (stimulus) and replace the tic with an alternative action instead of suppressing it.

The learning and execution of tics (and habits) in Tourette syndrome may be accelerated due to increased phasic dopamine bursts and dips onto cortico-striatal projections (Maia and Conceição, 2017; Conceição et al., 2017). At the same time, patients may have an increased propensity to execute such learned tics due to increased activation of the Go relative to No-Go pathway, as higher tonic dopamine levels increase and decrease the gain of D1 and D2 medium spiny neurons, respectively (Maia and Conceição, 2017; Conceição et al., 2017). However, according to this framework, both tic learning and tic execution take place in the dorsolateral (motor) loop through the putamen without the involvement of multiple loops or shortcuts. This view assumes the traditional perspective of parallel loops that localizes habitual actions in the sensorimotor dorsolateral loop (Yin and Knowlton, 2006). This organization however has been recently challenged by models which consider recent experiments regarding cortico-striatal projections (Baladron and Hamker, 2020; Balleine et al., 2015; Collins and Frank, 2013). A unique feature of our approach is the hierarchical organization of the multiple cortico-basal ganglia-thalamo-cortical loops.

Another line of evidence that has to be taken into account when modeling habitual behavior, comes from animal models: habit learning does not only involve the basal ganglia, but also critically depends on cortical areas. For instance, rats with lesions of the infralimbic cortex are unable to develop habits (Killcross and Coutureau, 2003). Further, the execution of established habits can be prevented if the same area is inactivated after learning (Smith et al., 2012; Coutureau and Killcross, 2003). Thus, the medial prefrontal cortex may be central to both learning and execution of habits. Our model explicitly includes a cortico-thalamo-cortical shortcut to model this dependence and suggests that enhanced habit formation in Tourette syndrome may be explained by increased speed of learning in this shortcut. Given the analogy of tics and habits, we encourage future models to include such shortcuts in order to investigate their dysfunction as a potential pathophysiological feature that contributes to the learning and execution of tics, and not just habits, in Tourette syndrome.

Around half of Tourette patients also present obsessive compulsive disorder (Goodman et al., 2006). Young patients with comorbid Tourette syndrom and obsessive-compulsive behavior have more severe tics (Lebowitz et al., 2012) and rely more on habitual behavior (Gillan et al., 2014). Further, current studies relate obsessive-compulsive disorder with a disruption in the balance between goal-directed behavior and habits (Gillan et al., 2011). According to our model, all these symptoms could be associated to shortcut malfunctions. A tentative compromise may be implemented by a two-step model of Tourette, where tics initially emerge by reduced levels of inhibition onto striatal projection neurons and then become manifested by enhanced habit formation.

### Role of Dopamine

The role of dopamine further supports the putative link between habits and tics. On the one hand, dopaminergic disturbances present a central suspected pathophysiological feature of Tourette syndrome (Buse et al., 2013; Maia and Conceição, 2017, 2018). Mice with excessive striatal dopamine show frequent rigid and complex action patterns and serve as an animal model of the Tourette syndrome (Berridge et al., 2005). On the other hand, dopamine takes a critical role during learning of habitual behaviors, although its influence on the execution of learned habits may diminish with growing cortical control (Ashby et al., 2010).

Our results suggest that excessive dopamine may increase habitual responses through an enhancement of the shortcut’s feedback to the striatum. Accordingly, it has been recently shown that rats develop habitual responses faster when they were exposed to the dopamine precursor levodopa (Gibson et al., 2020). Accelerated habit formation has also been observed in rats whose dopamine levels were increased through amphetamine sensitization (Nelson and Killcross, 2006). The inability of animals to form habits following lesions of the nigrostriatal dopamine system (Faure et al., 2005), the dorsolateral striatum (Yin et al., 2004), or the infralimbic cortex (Killcross and Coutureau, 2003; Coutureau and Killcross, 2003) reveals critical brain regions for habit formation and its dependence on dopamine. Experiments on rats further show that behavior becomes less dependent on dopamine with extended training (Choi et al., 2005), which could correspond to control being transferred from the loops to the shortcut. Indeed, it has been hypothesized that dopamine only affects the early learning of habits (Ashby et al., 2007). Assuming a link between tics and habits, dopamine-modulating medication may thus be more effective in preventing the learning of new tics instead of suppressing existing tics.

### Tourette Treatments May Affect Shortcut Connections

In our model with multiple loops, habitual behavior does not emerge if plasticity in the shortcut projections is disabled. Habits are not released from the dorsolateral loop alone, because thalamic cells are biased by a fast transmission of visual inputs via cortico-thalamic shortcut projections. The association between the respective cortical and thalamic cells slowly acquires over repeated trials, with the basal ganglia providing a teaching signal for the shortcut. Our simulation results suggest that this slow incremental learning process may be accelerated in the case of Tourette syndrome, benefiting the fast development and consolidation of habits which can manifest as tics. The effectiveness of habit reversal training (Dutta and Cavanna, 2013) and comprehensive behavioral intervention (Petruo et al., 2020) in treating tics may be explained by a rewiring in these shortcut projections. The initially learned maladaptive behavior (tic) can be replaced by another action if the connection pattern between cortical and thalamic cells in the shortcut can be modified.

A common target for deep brain stimulation in Tourette patients is the thalamic centromedian-parafascicular (CM-Pf) region (Mink, 2006; Britoa et al., 2019). According to our approach, the thalamus is a critical element of the shortcut, and indeed Tourette patients had increased basal ganglia-cortical and thalamo-cortical connectivity in a recent fMRI study (Ramkiran et al., 2019). Stimulation of the thalamus could therefore interfere with the spread of information through the cortico-thalamo-cortical pathway.

### Limitations and Open Issues

Although our model could recreate the task behavior of patients observed by Delorme et al. (2016), we did not include any explicit representation of tics, the core symptom of the Tourette syndrome. We already discussed the close resemblance of habits and tics (Leckman and Riddle, 2000). Our model however could be complemented with a direct representation of tics in order to simulate a wider range of experiments. For instance, Caligiore et al. (2017) proposed a neuro-computational model of a single loop in order to explain tic generation in a pharmacologic monkey model of motor tics studied by McCairn et al. (2013). According to their model, tics are generated as a consequence of dysfunctional interaction between the cortico-basal-ganglia loops and the cerebellum. In their simulations, enhanced phasic bursts of dopamine made the basal ganglia overly sensitive to cortical noise, producing undesired activation of the premotor cortex which is understood as tic initiation. Although their model uses similar firing rate units and has a comparable structure to ours, it does not include plasticity and can therefore not learn to solve any task.

A central finding in support of the hypothesis of reduced striatal inhibition is the loss of interneurons observed in stereological analyses of post-mortem brains of Tourette patients (Kalanithi et al., 2005; Kataoka et al., 2010). As our model of the striatum is composed only of medium spiny neurons, we have approximated the loss of inhibitory interneurons by a reduction of local inhibitory connections. The loss of inhibitory interneurons may result in a more complex change than approximated in the present model version.

Delorme et al. (2016) not only reported the behavioral results replicated here, but also used diffusion tensor imaging to study the structural connectivity within the basal ganglia. They found that a higher amount of responses to devaluated outcomes was correlated with an increase in the connectivity of the motor network. Possible effects of shortcuts however were excluded from their analysis as only the posterior sensorimotor putamen and the anterior caudate nucleus were used as seeds. Based on our results we suggest to further include cortical or thalamic areas as seed regions in future experiments.

Our modeling results suggest that shortcut connections are crucial to learn and engage in habitual behavior. While our understanding of the dependence of habit learning on cortical areas such as the infralimbic cortex in rats is growing (Killcross and Coutureau, 2003; Coutureau and Killcross, 2003; Smith et al., 2012), a comparable site in the human brain is yet to be found. Our model predicts the development of habitual behavior on basis of active shortcut connections. Both pathophysiological model alterations, disturbing tonic dopamine signaling or reducing local inhibition in the striatum, indirectly modulate shortcut activity and lead to increased rates of responses to stimuli associated with devalued outcomes in our model. Despite the uncertainties about the cortical sites mediating shortcuts in humans, our model inspires novel research ideas regarding the conceptual understanding of tics as habits and the search for the respective neural underpinning in corticothalamic shortcut connections, which may lead to new treating options for patients with Tourette syndrome.

As a final remark, our model has been originally developed to explain habit formation in animals (Baladron and Hamker, 2020). Habits in Delorme et al. (2016) however, refer rather to a more cognitive outcome-insensitive behavioral control. Humans appear in general more sensitive to outcome-devaluation and thus, less sensitive to habits (de Wit et al., 2018). Despite these discrepancies between human and animal studies of habit formation, our model may rather help to understand such differences, as habits imposed by the shortcut could be diminished by cognitive control.

## Methods

### Model details

The cortex - basal ganglia model is based on the one proposed by Baladron and Hamker (2020), which further extends that of Schroll et al. (2014) to include multiple loops. Each basal ganglia loop includes the direct pathway, through the striatum and the internal globus pallidus (GPi), the short indirect pathway, through the striatum and the external globus pallidus (GPe), and the hyperdirect pathway through the STN (Figure 2B).

Neural populations modeled by a number of rate coded neurons which follow the differential equation:

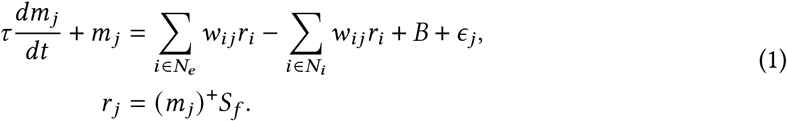

where *m_j_* is the membrane potential of cell *j*, *r_j_* the firing rate of cell *j*, *τ* a time constant, *w_ij_* the weight between presynaptic cell *i* and postsynaptic cell *j*, *N_e_* are the cells with an excitatory synapse to cell *j*, *N_i_* are the cells with an inhibitory synapse to cell *j*, *B* is a baseline activity and *ϵ_j_* is a noise term drawn from a uniform distribution, ()^+^ converts negative numbers to 0. *S_f_* is a scaling factor used to simulate disturbed altered response modulation. On control models it is set to 1.

The striatum of each loop is divided into two main groups: One for D1 dopamine receptor expressing cells, and one for D2 dopamine receptor cells. Each group contains an additional smaller set of cells that implement cortical feedback (Figure 2A). These neurons have afferent connections only from the cortical cells of their corresponding loop and project to both the GPi and the GPe. A similar mechanism was used in the original model (Schroll et al., 2014) to implement thalamic feedback and facilitate learning in each pathway.

Learning in the cortico-striatal projections follows a dopamine-modulated covariance learning rule:

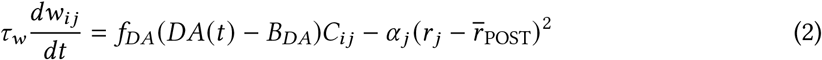

where *w_ij_* is the weight of the synapse between presynaptic cell *i* and postsynaptic cell *j*, *f_DA_* implements dopamine modulation and it is a function of the difference between the current dopamine level *DA(t)* and the baseline dopamine level *B_DA_*, *C_ij_* is a correlation measure and 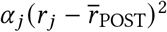 a normalization term that limits the weight growth.

The function *f_DA_* ensures that a phasic dopamine increase enhances long-term potentiation in D1 receptor expressing cells and long-term depression in D2 receptor expressing cells (Shen et al., 2008; Villagrasa et al., 2018). The expression for this function is:

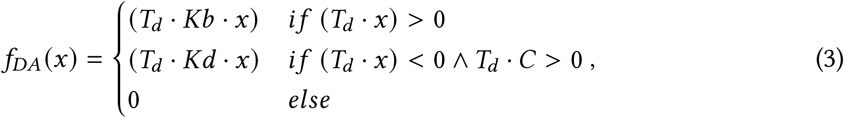

where *T_d_* controls the effect of dopamine, *Kb* and *Kd* the size of increases and decreases of the weights. *T_d_* is 1 on projections to D1 expressing cells and −1 to D2 expressing cells.

The correlation measure *C_ij_* is defined such that only connections between active cells are subject to plasticity. In the dorsomedial loop, it follows:

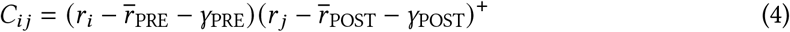

where *r_i_* and *r_j_* are the rate of the presynaptic and postsynaptic cells, 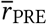 and 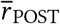 are the mean firing rate of the presynaptic and postsynaptic populations and *γ*_PRE_ and *γ*_POST_ are thresholds.

The correlation measure is different in the dorsolateral striatum, where a trace (*Tr_j_*) is required. The trace allows plasticity to occur only between cells that were active during the period in which an action was selected. Active cells during this period may be different to those active once dopamine increases or decreases due to the integration of environmental information done by cortical cells. Correlation is then defined as:

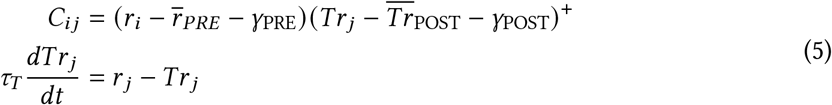

The normalization term in equation 2 is only active if the activity of the postsynaptic cell is larger than a fixed threshold (*m*^MAX^). This is controlled through *α* which is governed by the following differential equation:

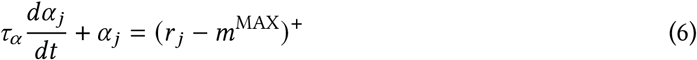

The model also includes plasticity in the striato-pallidal connections. Different than corticostriatal connections, the learning rule in these projections ensures that plasticity occurs only when the activity of the pallidal cell is below the mean. This is required as for the selection of an objection/action a decrease is required and not an increase in the firing rate is rquired. The weight change then follows:

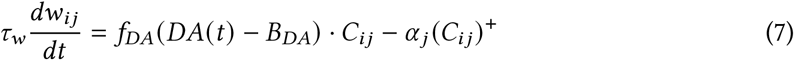

with *f_DA_* (*x*) and *a_j_* defined in equation 3 and with:

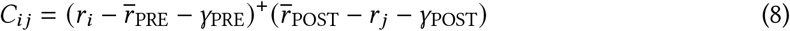

for the projections in the dorsomedial loop and:

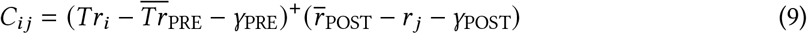

for the projections in the dorsolateral loop.

Cortical projections to the subthalamic nucleus also follow equation 2 with *T_d_* = 1. Projections from the subthalamic nucleus to the pallidum follow equation 7 with *T_d_* = 1.

A different set of dopaminergic cells project to each loop. Each dopaminergic cell in the dorsomedial loop is associated with one of the six possible outcomes. In the dorsolateral loop, each dopaminergic cell is associated with one possible button. On both loops, dopaminergic cells project to all nuclei. Their activation is governed by the following equation:

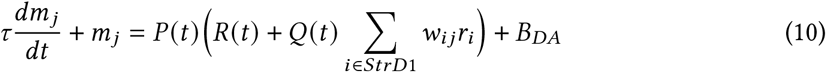

where *B* represents the baseline dopamine level at resting conditions. The function *P(t)* controls the timing of phasic changes. This function is 1 after every action and 0 otherwise. The function *R(t)* controls the size of the phasic changes and it is different in both loops. In the dorsomedial loop, *R(t)* is (1 − *B*) if the outcome associated with the cell is obtained and 0 otherwise. In the dorsolateral loop, *R(t)* is (1 − *B*) only if the button associated with the cell is pressed. Following the reward prediction error hypothesis, the response of dopaminergic cells is reduced through inhibitory plastic connections from the striatal D1 cells of the corresponding loop. Additionally, the function *Q(t)* is used to recreate the strong drop observed when a reward is predicted but not obtained. In the dorsomedial loop, *Q(t)* = −10 in unrewarded trials and *Q(t)* = −1 in rewarded trials.

Although dopamine-dependent plasticity is implemented in both loops, the respective dopamine signals differ and provide each loop with the appropriate prediction error information (Engelhard et al., 2019; Rusu and Pennartz, 2020; Poulin et al., 2018). As in Baladron and Hamker (2020), dopamine in the dorsomedial loop encodes a classical reward prediction error signal while in the dorsolateral loop, it encodes an action consequence prediction error. Ongoing learning leads to a selective increase of the inhibitory projection from the striatum to the dopaminergic cells (prediction part) and thus reduces the error.

Successful trials increase the weights of the inhibitory connections from the striatal cells to the dopaminergic cells, following the equation:

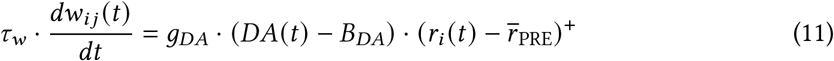

where *w_ij_* is the weight between the striatal cell *i* and the dopaminergic cell *j*,

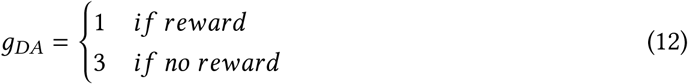

for the dorsomedial cells and *g_DA_* = 1 for the dorsolateral cells, DA(t) is the sum of dopaminergic inputs to the striatal cell *i*, *r_i_* is the firing rate of striatal cell *i*, 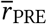 is the mean of the presynaptic layer and *τ_w_* is a time constant (3,000 ms for the dorsomedial loop and 12,000 ms for the dorsolateral loop).

Plasticity in the shortcut is not modulated by dopamine. These connections follow the correlation learning rule:

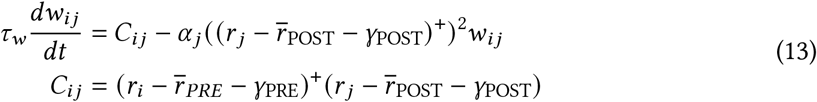

Further, these connections present slower changes than the cortico-striatal projections (controlled through different values of *τ_w_*). A pattern can only be obtained if the basal ganglia select the same action multiple times.

All differential equations are numerically solved using the Euler method with a time step of 1 ms using the neuro-simulator ANNarchy version 4.6 (Vitay et al., 2015).

### Task mapping

In order to simulate the task of Delorme et al. (2016) we included a set of 12 cells encoding the visual stimuli, 2 encoding each of the possible 6 fruit icons outside the boxes. These input cells had plastic connections to the dorsomedial loop (striatum and subthalamic nucleus) and to the shortcut (Figure 2B).

An additional set of 30 input cells encoded a desire for a possible outcome fruit. Each of the 6 possible outcome fruits was therefore encoded by 5 cells. Fixed connections were included from these cells to the striatum and the subthalamic nucleus of the dorsomedial loop, both which were further divided into 6 groups of 4 neurons each, each receiving excitatory input from a different outcome fruit.

The cortex, GPi and GPe of each loop contain 2 cells each. In the dorsomedial loop these cells represent the possible keys that the subject may press while in the dorsolateral loop they represent the necessary hand movement to reach the buttons. Cortical cells in the dorsomedial loop receive an external input that increases their firing rate if their corresponding key was pressed at the end of a trial (independent of the selection done by the loop).

At the beginning of each simulated trial, the firing rate of the visual stimuli cells encoding one fruit was set to 0.5 and the firing rate of the cells encoding the corresponding outcome fruit was set to 1.0. Then, the network was simulated until either 600ms have elapsed or a cell in the cortex of the dorsolateral loop reached a threshold of 0.2. When the model did not reach the threshold, it was considered as if no response was produced. On trials in which the threshold was reached, the final decision was computed stochastically according to the categorical probability distribution obtained using the activity of the cortical cells in the dorsolateral loop:

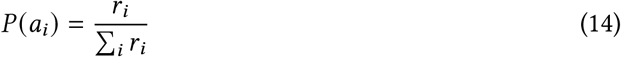

where *P (a_i_*) is the probability of selecting the action associated with the cortical cell *i* and *r_i_* is the firing rate of the cortical cell *i*.

After the decision, the phasic level of dopamine was changed according to equation 10 for an additional 100ms of simulation. The dopamine modulated learning rule was therefore only active during this period.

Finally, the input was removed and an inter-trial period of 700ms was simulated. This allowed the network to reach a stable initial condition before the next trial.

Each model was trained for 10 blocks in which each possible stimulus was presented twice. For each of the conditions described in the results section, 50 models were simulated, each with different initial conditions and noise values. After the training period, outcome devaluation and stimulus devaluation experiments were simulated. On outcome devaluation trials, no outcome encoding cell was activated. On stimulus devaluation trials, the firing rate of the corresponding visual input cells was set to a random value sampled from a normal distribution with mean 0.28 and standard deviation 0.05. This represents that the stimuli was still seen, but was unattended. The response rate in each case was computed as the number of trials in which the threshold was reached, divided by the total number.

Disturbed tonic dopamine signaling was simulated by multiplying the firing rate (*r* in Equation of the striatal D1 neurons by a factor bigger than 1 and the firing rate of striatal D2 neurons by a factor between 0 and 1. Disturbed phasic dopamine was simulated by decreasing the baseline value (*B_DA_*). Such a change enhances phasic learning, as the plasticity rule depends on the difference between the current dopamine level and the baseline (*f_DA_ DA(t) − B_DA_*) and is only applied after the response. Reduced striatal inhibition was simulated by multiplying the weight of the local connection between striatal cells by different factors, as reported in the results section.

### Statistical analysis

Statistical significance between model groups was measured using a permutation test. On each test we first pooled both groups together and then re-sampled into two groups 1,000,000 times. For each sample, we computed the difference between the two groups and then used this result to estimate the p-value as the proportion of samples that had a difference greater or equal than the original value. If the estimated p-value was smaller than 0.005, the difference was considered significant.

## Acknowledgments

We thank Izhar Bar-Gad, Jonathan Rubin, Christian Beste, Lieneke Janssen and Kathleen Wiencke for their helpful comments on previous versions of this manuscript. This work was supported by the Federal Ministry of Education and Research grant “Multilevel neuro-computational models of basal ganglia dysfunction in Tourette syndrome” (BMBF 01GQ1707) as part of the program “CRCNS US-German-Israeli collaboration on computational neuroscience” jointly with Izhar Bar-Gad and Jonathan Rubin. C.S. was further supported by BMBF and the Max Planck Society.

## References

Adams, C. D. (1982). Variations in the sensitivity of instrumental responding to reinforcer devaluation. The Quarterly Journal of Experimental Psychology Section B, 34(2b):77–98.

Ashby, F. G., Ennis, J. M., and Spiering, B. J. (2007). A neurobiological theory of automaticity in perceptual categorization. Psychological review, 114(3):632.

Ashby, F. G., Turner, B. O., and Horvitz, J. C. (2010). Cortical and basal ganglia contributions to habit learning and automaticity. Trends in cognitive sciences, 14(5):208–215.

Assous, M. and Tepper, J. M. (2019). Excitatory extrinsic afferents to striatal interneurons and interactions with striatal microcircuitry. European Journal of Neuroscience, 49(5):593–603.

Baladron, J. and Hamker, F. H. (2015). A spiking neural network based on the basal ganglia functional anatomy. Neural Networks, 24:1–13.

Baladron, J. and Hamker, F. H. (2020). Habit learning in hierarchical cortex-basal ganglia loops. European Journal of Neuroscience.

Balleine, B. W., Dezfouli, A., Ito, M., and Doya, K. (2015). Hierarchical control of goal-directed action in the cortical–basal ganglia network. Current Opinion in Behavioral Sciences, 5:1–7.

Balleine, B. W., Killcross, A. S., and Dickinson, A. (2003). The effect of lesions of the basolateral amygdala on instrumental conditioning. Journal of Neuroscience, 23:666–675.

Berridge, K. C., Aldridge, J. W., Houchard, K. R., and Zhuang, X. (2005). Sequential super-stereotypy of an instinctive fixed action pattern in hyper-dopaminergic mutant mice: a model of obsessive compulsive disorder and Tourette’s. BMC biology, 3(1):4.

Beste, C. and Münchau, A. (2018). Tics and Tourette syndrome—surplus of actions rather than disorder? Movement Disorders, 33(2):238–242.

Brand, N., Geenen, R., Oudenhoven, M., Lindenborn, B., van der Ree, A., Cohen-Kettenis, P., and Buitelaar, J. K. (2002). Brief report: Cognitive functioning in children with Tourette’s syndrome with and without comorbid ADHD. Journal of Pediatric Psychology, 27:203–208.

Brandt, V. C., Beck, C., Sajin, V., Baaske, M. K., Bäumer, T., Beste, C., Anders, S., and Münchau, A. (2016). Temporal relationship between premonitory urges and tics in Gilles de la Tourette syndrome. Cortex, 77:24–37.

Britoa, M., Teixeira, M. J., Mendes, M. M., Françac, C., Iglesio, R., Barbosac, E. R., and Cury, R. G. (2019). Exploring the clinical outcomes after deep brain stimulation in tourette syndrome. Journal of the Neurological Sciences, 402:48–51.

Bronfeld, M. and Bar-Gad, I. (2013). Tic disorders: what happens in the basal ganglia? The Neuroscientist, 19(1):101–108.

Buse, J., Schoenefeld, K., Münchau, A., and Roessner, V. (2013). Neuromodulation in Tourette syndrome: dopamine and beyond. Neuroscience & Biobehavioral Reviews, 37(6):1069–1084.

Caligiore, D., Mannella, F., Arbib, M. A., and Baldassarre, G. (2017). Dysfunctions of the basal ganglia-cerebellar-thalamo-cortical system produce motor tics in Tourette syndrome. Plos Computational Biology, 13:1–34.

Capriotti, M. R., Brandt, B. C., Turkel, J. E., Lee, H.-J., and Woods, D. W. (2014). Negative reinforcement and premonitory urges in youth with Tourette syndrome: an experimental evaluation. Behavior modification, 38(2):276–296.

Choi, W. Y., Balsam, P. D., and Horvitz, J. C. (2005). Extended habit training reduces dopamine mediation of appetitive response expression. Journal of Neuroscience, 20:6729–6733.

Collins, A. G. and Frank, M. J. (2013). Cognitive control over learning: creating, clustering and generalizing task-set structure. Psychological Review, 120:190–229.

Conceição, V. A., Ângelo Dias, Farinha, A. C., and Maia, T. V. (2017). Premonitory urges and tics in Tourette syndrome: computational mechanisms and neural correlates. Current opinion in neurobiology, 46:187–199.

Corbit, L. H., Muir, J. L., and Balleine, B. W. (2001). The role of the nucleus accumbens in instrumental conditioning: Evidence of a functional dissociation between accumbens core and shell. Journal of Neuroscience, 21:3251–3260.

Coutureau, E. and Killcross, S. (2003). Inactivation of the infralimbic prefrontal cortex reinstates goal-directed responding in overtrained rats. Behavioural brain research, 146(1-2):167–174.

de Wit, S., Kindt, M., Knot, S. L., Verhoeven, A. A. C., Robbins, T. W., Gasull-Camos, J., Evans, M., Mirza, H., and Gilla, C. M. (2018). Shifting the balance between goals and habits: Five failures in experimental habit induction. Journal of Experimental Psychology General, 147:1043–1065.

Delorme, C., Salvador, A., Valabregue, R., Roze, E., Palminteri, S., Vidailhet, M., de Wit, S., Robbins, T., Hartmann, A., and Worbe, Y. (2016). Enhanced habit formation in Gilles de la Tourette syndrome. Brain, 139(2):605–615.

Dutta, N. and Cavanna, A. E. (2013). The effectiveness of habit reversal therapy in the treatment of Tourette syndrome and other chronic tic disorders: a systematic review. Functional neurology, 28(1):7.

Eddy, C. M. and Cavanna, A. E. (2013). Altered social cognition in Tourette syndrome: nature and implications. Behavioral Neurology, 27:15–22.

Engelhard, B., Finkelstein, J., Cox, J., Fleming, W., Jang, H. J., Ornelas, S., Koay, S. A., Thiberge, S. Y., Daw, N. D., Tank, D. W., et al. (2019). Specialized coding of sensory, motor and cognitive variables in VTA dopamine neurons. Nature, 570(7762):509–513.

Faure, A., Haberland, U., Condé, F., and El Massioui, N. (2005). Lesion to the nigrostriatal dopamine system disrupts stimulus-response habit formation. Journal of Neuroscience, 25(11):2771–2780.

Fisher, S. D., Robertson, P. B., Black, M. J., Redgrave, P., Sagar, M. A., Abraham, W. C., and Reynolds, J. N. (2017). Reinforcement determines the timing dependence of corticostriatal synaptic plasticity in vivo. Nature Communications, 8:1–13.

Gerfen, C. R. and Surmeier, D. J. (2011). Modulation of striatal projection systems by dopamine. Annual Review of Neuroscience, 34:441–466.

Gibson, A. S., Keefe, K. A., and Furlong, T. M. (2020). Accelerated habitual learning resulting from L-dopa exposure in rats is prevented by N-acetylcysteine. Pharmacology Biochemistry and Behavior, 198:1730–1733.

Gillan, C. M., Morein-Zamir, S., Urcelay, G. P., Sule, A., Voon, V., Apergis-Schoute, A. M., Fineberg, N. A., Sahakian, B. J., and Robbins, T. W. (2014). Enhanced avoidance habits in obsessive-compulsive disorder. Biological Psychiatry, 75:631–638.

Gillan, C. M., Papmeyer, M., Morein-Zamir, S., Sahakian, B. J., Fineberg, N. A., and Trevor W Robbins, S. d. W. (2011). Disruption in the balance between goal-directed behavior and habit learning in obsessive-compulsive disorder. The American Journal of Psychiatry, 168:718–726.

Gönner, L., Vitay, J., and Hamker, F. H. (2017). Predictive place-cell sequences for goal-finding emerge from goal memory and the cognitive map: a computational model. Frontiers in computational neuroscience, 11:1–19.

Goodman, W. K., Storch, E. A., Geffken, G. R., and Murphy, T. K. (2006). Obsessive-compulsive disorder in Tourette syndrome. Journal of Child Neurology, 21:704–714.

Groenewegen, H. J., Wright, C. I., Beijer, A. V., and Voorn, P. (1999). Convergence and segregation of ventral striatal inputs and outputs. Annals of the New York academy of sciences, 877:49–63.

Groenewegen, H. J., Wright, C. I., and Uylings., H. B. (1997). The anatomical relationships of the prefrontal cortex with limbic structures and the basal ganglia. Journal of Psychopharmacology, 11:99–106.

Haber, S. N. (2016). Corticostriatal circuitry. Dialogues in clinical neuroscience, 18(1):7.

Kalanithi, P. S., Zheng, W., Kataoka, Y., DiFiglia, M., Grantz, H., Saper, C. B., Schwartz, M. L., Leckman, J. F., and Vaccarino, F. M. (2005). Altered parvalbumin-positive neuron distribution in basal ganglia of individuals with Tourette syndrome. Proceedings of the National Academy of Sciences, 102(37):13307–13312.

Kataoka, Y., Kalanithi, P. S., Grantz, H., Schwartz, M. L., Saper, C., Leckman, J. F., and Vaccarino, F. M. (2010). Decreased number of parvalbumin and cholinergic interneurons in the striatum of individuals with Tourette syndrome. Journal of Comparative Neurology, 518(3):277–291.

Killcross, S. and Coutureau, E. (2003). Coordination of actions and habits in the medial prefrontal cortex of rats. Cerebral cortex, 13(4):400–408.

Kleimaker, M., Takacs, A., Conte, G., Onken, R., Verrel, J., Bäumer, T., Münchau, A., and Beste, C. (2020). Increased perception-action binding in Tourette syndrome. Brain, 143:1934–1945.

Kwak, C., Dat Vuong, K., and Jankovic, J. (2003). Premonitory sensory phenomenon in Tourette’s syndrome. Movement disorders, 18(12):1530–1533.

Lebowitz, E. R., Motlagh, M. G., Katsovich, L., King, R. A., Lombroso, P. J., Grantz, H., Lin, H., Bentley, M. J., Gilbert, D. L., Singer, H. S., Coffey, B. J., the Tourette Syndrome Study Group, Kurlan, R. M., and Leckman, J. F. (2012). Tourette syndrome in youth with and without obsessive compulsive disorder and attention deficit hyperactivity disorder. European Child & Adolescent Psychiatry, 21:451–457.

Leckman, J. F. and Riddle, M. A. (2000). Tourette’s syndrome: When habit-forming systems form habits of their own? Neuron, 28:349–354.

Leckman, J. F., Walker, D. E., and Cohen, D. J. (1993). Premonitory urges in Tourette’s syndrome. The American journal of psychiatry.

Maia, T. V. and Conceição, V. A. (2017). The roles of phasic and tonic dopamine in tic learning and expression. Biological psychiatry, 82(6):401–412.

Maia, T. V. and Conceição, V. A. (2018). Dopaminergic disturbances in Tourette syndrome: an integrative account. Biological psychiatry, 84(5):332–344.

Maia, T. V. and Conceição, V. A. (2017). The roles of phasic and tonic dopamine in tic learning and expression. Biological Psychiatry, 82:401–412.

McCairn, K., Iriki, A., and Isoda, M. (2013). Global dysrhythmia of cerebro-basal ganglia-cerebellar networks underlies motor tics following striatal disinhibition. Journal of Neuroscience, 33:697–708.

McCairn, K. W., Bronfeld, M., Belelovsky, K., and Bar-Gad, I. (2009). The neurophysiological correlates of motor tics following focal striatal disinhibition. Brain, 132(8):2125–2138.

Mink, J. W. (2006). Clinical review of DBS for Tourette syndrome. Frontiers in Bioscience, E1:72–76.

Nelson, A. and Killcross, S. (2006). Amphetamine exposure enhances habit formation. Journal of Neuroscience, 26(14):3805–3812.

Openneer, T. J., Tárnok, Z., Bognar, E., Benaroya-Milshtein, N., Garcia-Delgar, B., Morer, A., Steinberg, T., Hoekstra, P. J., and Dietrich, A. (2019). The premonitory urge for tics scale in a large sample of children and adolescents: psychometric properties in a developmental context. an emtics study. European Child & Adolescent Psychiatry, pages 1–14.

Palminteri, S., Lebreton, M., Worbe, Y., Grabli, D., Hartmann, A., and Pessiglione, M. (2009). Pharmacological modulation of subliminal learning in Parkinson’s and Tourette’s syndromes. Proceedings of the National Academy of Sciences, 106(45):19179–19184.

Palminteri, S., Lebreton, M., Worbe, Y., Hartmann, A., Lehéricy, S., Vidailhet, M., Grabli, D., and Pessiglione, M. (2011). Dopamine-dependent reinforcement of motor skill learning: evidence from Gilles de la Tourette syndrome. Brain, 134(8):2287–2301.

Peterson, B. S., Skudlarski, P., Anderson, A. W., Zhang, H., Gatenby, J. C., Lacadie, C. M., Leckman, J. F., and Gore, J. C. (1998). A functional magnetic resonance imaging study of tic suppression in Tourette syndrome. Archives of general psychiatry, 55(4):326–333.

Peterson, B. S., Thomas, P., Kane, M. J., Scahill, L., Zhang, H., Bronen, R., King, R. A., Leckman, J. F., and Staib, L. (2003). Basal ganglia volumes in patients with Gilles de la Tourette syndrome. Archives of general psychiatry, 60(4):415–424.

Petruo, V., Bodmer, B., Bluschke, A., Münchau, A., Roessner, V., and Beste, C. (2020). Comprehensive behavioral intervention for tics reduces perception-action binding during inhibitory control in Gilles de la Tourette syndrome. Nature Scientific Reports, 10:1–8.

Pogorelov, V., Xu, M., Smith, H. R., Buchanan, G. F., and Pittenger, C. (2015). Corticostriatal interactions in the generation of tic-like behaviors after local striatal disinhibition. Experimental neurology, 265:122–128.

Poulin, J.-F., Caroni, G., Hofer, C., Cui, Q., Helm, B., Ramakrishnan, C., Chan, C. S., Dombeck, D., Deisseroth3, K., and Awatramani1, R. (2018). Mapping projections of molecularly defined dopamine neuron subtypes using intersectional genetic approaches. Nature Neuroscience, 21:1260–1271.

Puts, N. A. J., Harris, A. D., Crocetti, D., Nettles, C., Singer, H. S., Tommerdahl, M., Edden, R. A. E., and Mostofsky, S. H. (2015). Reduced GABAergic inhibition and abnormal sensory symptoms in children with Tourette syndrome. Journal of Neurophysioloy, 114:808–817.

Ramkiran, S., Heidemeyer, L., Gaebler, A., Shah, N. J., and Neuner, I. (2019). Alterations in basal ganglia-cerebello-thalamo-cortical connectivity and whole brain functional network topology in Tourette’s syndrome. NeuroImage: Clinical, 24:101998.

Redgrave, P., Rodriguez, M., Smith, Y., Rodriguez-Oroz, M. C., Lehericy, S., Bergman, H., Agid, Y., DeLong, M. R., and Obeso, J. A. (2010). Goal-directed and habitual control in the basal ganglia: implications for Parkinson’s disease. Nature Reviews Neuroscience, 11:760–7720.

Rusu, S. I. and Pennartz, C. M. A. (2020). Learning, memory and consolidation mechanisms for behavioral control in hierarchically organized cortico-basal ganglia systems. Hippocampus, 64:73–98.

Schroll, H., Vitay, J., and Hamker, F. H. (2014). Dysfunctional and compensatory synaptic plasticity in parkinsons disease. European Journal of Neuroscience, 39.

Shen, W., Flajolet, M., Greengard, P., and Surmeier, D. J. (2008). Dichotomous dopaminergic control of striatal synaptic plasticity. Science, 321(5890):848–851.

Shephard, E., Groom, M. J., and Jackson, G. M. (2019). Implicit sequence learning in young people with Tourette syndrome with and without co-occurring attention-deficit/hyperactivity disorder. Journal of Neuropsychology, 13(3):529–549.

Sherman, S. M. and Guillery, R. W. (2011). Distinct functions for direct and transthalamic cortico-cortical connections. J Neurophysiol, 106:1068–1077.

Singer, H. S., Butler, I. J., Tune, L. E., Seifert Jr, W. E., and Coyle, J. T. (1982). Dopaminergic dysfunction in Tourette syndrome. Annals of Neurology: Official Journal of the American Neurological Association and the Child Neurology Society, 12(4):361–366.

Singer, H. S., Szymanski, S., Giuliano, J., Yokoi, F., Dogan, A. S., Brasic, J. R., Zhou, Y., Grace, A. A., and Wong, D. F. (2002). Elevated intrasynaptic dopamine release in Tourette’s syndrome measured by PET. American Journal of Psychiatry, 159(8):1329–1336.

Smith, K. S. and Graybiel, A. M. (2013). A dual operator view of habitual behavior reflecting cortical and striatal dynamics. Neuron, 79:361–374.

Smith, K. S., Virkud, A., Deisseroth, K., and Graybiel, A. M. (2012). Reversible online control of habitual behavior by optogenetic perturbation of medial prefrontal cortex. Proceedings of the National Academy of Sciences, 109(46):18932–18937.

Surmeier, D. J., Ding, J., Day, M., Wang, Z., and Shen, W. (2007). D1 and D2 dopamine-receptor modulation of striatal glutamatergic signaling in striatal medium spiny neurons. Trends in Neuroscience, 30:228–235.

Villagrasa, F., Baladron, J., Vitay, J., Schroll, H., Antzoulatos, E. G., Miller, E. K., and Hamker, F. H. (2018). On the role of cortex-basal ganglia interactions for category learning: A neurocomputational approach. Journal of Neuroscience, 31:9551–9562.

Vinner, E., Israelashvili, M., and Bar-Gad, I. (2017). Prolonged striatal disinhibition as a chronic animal model of tic disorders. Journal of neuroscience methods, 292:20–29.

Vitay, J., Dinkelbach, H. U., and Hamker, F. H. (2015). ANNarchy: a code generation approach to neural simulations on parallel hardware. Frontiers in Neuroinformatics, 9:1–10.

Wickens, J. R. (2009). Synaptic plasticity in the basal ganglia. Behavioural Brain Research, 199:119–128.

Wong, D. F., Brašić, J. R., Singer, H. S., Schretlen, D. J., Kuwabara, H., Zhou, Y., Nandi, A., Maris, M. A., Alexander, M., Ye, W., et al. (2008). Mechanisms of dopaminergic and serotonergic neurotransmission in Tourette syndrome: clues from an in vivo neurochemistry study with pet. Neuropsychopharmacology, 33(6):1239–1251.

Yael, D., Tahary, O., Gurovich, B., Belelovsky, K., and Bar-Gad, I. (2019). Disinhibition of the nucleus accumbens leads to macro-scale hyperactivity consisting of micro-scale behavioral segments encoded by striatal activity. Journal of Neuroscience, 39:5897–5909.

Yin, H. H. (2017). The basal ganglia in action. The Neuroscientist, 23(3):299–313.

Yin, H. H. and Knowlton, B. J. (2006). The role of the basal ganglia in habit formation. Nature Reviews Neuroscience, 7(6):464–476.

Yin, H. H., Knowlton, B. J., and Balleine, B. W. (2004). Lesions of dorsolateral striatum preserve outcome expectancy but disrupt habit formation in instrumental learning. European journal of neuroscience, 19(1):181–189.

